# Mature Citrus Leaves Undergo Coordinated Photosynthetic Downregulation to Support Flush-Driven Carbon and Nitrogen Sink Demand

**DOI:** 10.64898/2026.03.09.710566

**Authors:** Syed Bilal Hussain, Qingyan Meng, Sheng-yang Li, Yu Wang, Christopher Vincent

**Affiliations:** Citrus Research & Education Center, University of Florida, Lake Alfred, FL, 33850, USA; School of Agriculture and Food Science, University College Dublin, Dublin 4, D04 V1W8, Ireland; Department of Horticulture, Michigan State University, East Lansing, MI 48824, USA

**Author notes:** **Date of submission: 3/9/2026**.

**Keywords:** Source□sink dynamics, Carbon export, Flush phenology, Leaf position, Photosynthetic regulation

## Abstract

The source□sink attenuation hypothesis suggests that plants regulate carbon fixation in response to fluctuations in sink demands. Many evergreen trees exhibit flushing growth patterns, where new shoot development generates a strong, transient demand for both carbon and nitrogen that may influence the function of mature leaves. This study examined the source–sink attenuation hypothesis in the context of vegetative sink growth by investigating the photosynthetic capacity and nitrogen dynamics in mature citrus leaves across three stages of flush development. In contrast to expectations, photosynthesis declined as flush growth progressed. Early flush initiation induced stomatal limitation in mature leaves, whereas as sink demand from further shoot growth continued carboxylation capacity and Rubisco abundance declined, despite relatively stable total leaf nitrogen. These results suggest that mature leaves undergo selective protein retooling under prolonged sink demand, constraining CO□ fixation while maintaining C export. Overall, this study revealed that under strong combined N and C sink demands, mature citrus leaves function primarily as regulated carbon conduits rather than dynamically upregulating photosynthesis, providing new insight into source–sink coordination in woody perennial species.

**Highlight:** Citrus flush growth shows that mature leaves suppress photosynthesis through stomatal and biochemical regulation while reallocating carbon and nitrogen to support new shoot development, challenging classic source–sink theory.

## Introduction

Perennial plants play a vital role in ecosystem and biosphere functions by contributing to carbon (C) sequestration, nutrient cycling, and soil health maintenance (Van Driesche *et al*., 2010; Werling *et al*., 2014). However, optimizing their productivity remains a challenge because of the complex interplay between CO_2_ fixation, growth phenology, and nutrient availability, among other environmental factors. In agriculture, where perennial crops are increasingly valued for both their ecological and economic benefits, understanding how photosynthetic carbon is produced and allocated across tissues is critical for improving resource use efficiency (Lamichhane and Alletto, 2022).

C and nitrogen (N) regulation in plants is primarily mediated by source□sink interactions (Lemoine *et al*., 2013; Paul and Foyer, 2001; Poorter *et al*., 2012). Mature leaves serve as the primary sources of assimilates, whereas developing organs, including young leaves, roots, seeds, and meristems, act as dynamic sinks (Millard and Grelet, 2010; Osorio *et al*., 2014; Smith *et al*., 2018; Xiao-Li *et al*., 2022). In evergreen perennials, repeated cycles of vegetative flushes create dynamic changes in sink strength, altering the demand for exported photoassimilates and potentially triggering feedback responses in source leaves (Chapin III *et al*., 1986; Richardson *et al*., 2013). Importantly, C fixation, N assimilation, and protein turnover are increasingly viewed as coordinated processes that plants dynamically adjust to balance growth, resource-use efficiency, and developmental demand, rather than maintaining maximal photosynthetic capacity at all times (Liu *et al*., 2025).

While numerous studies have addressed the effects of leaf age and position on photosynthetic capacity, the findings remain inconsistent (Gonzalez□Real and Baille, 2000; Henning *et al*., 1979; Kitajima *et al*., 2002). Some studies report a decline in photosynthesis with increasing leaf age due to reallocation of N or reduced sink strength (Crous *et al*., 2021; Menezes *et al*., 2022; Xie and Luo, 2003), whereas others observe sustained or even enhanced photosynthetic capacity in mature leaves (Frank, 1981; Proietti *et al*., 2000). Moreover, the role of leaf position in mediating source strength and carbon export remains underexplored.

Citrus trees display a growth habit that is useful in studying source□sink dynamics. Citrus trees exhibit a flushing phenology in which shoots emerge with all leaf primordia and complete their growth in approximately three weeks (Syvertsen, 1985). The pattern of phenology involves multiple cycles per year in which newly matured shoots are quiescent (3-4 weeks), followed by periods of rapid shoot growth (3 weeks), providing an ideal system for investigating C dynamics and resource allocation in perennials. The mature leaves subtending the new shoots are still in the early period of their lifespan, given that the next flush typically occurs at 1-2 months and that the citrus leaf lifespan is greater than 17 months in the field (Wallace *et al*., 1954).

Recent work by Li and Vincent (2022) demonstrated that the emergence of new shoots enhances carbon export from mature leaves, whereas work by Kriedemann (1970) showed that basal leaves tend to supply C to roots and stems (basipetal translocation), whereas apical leaves preferentially support new shoot growth (apical translocation). In the case in which roots are sinks, Li et al. (2024) reported increased photosynthesis in response to defoliation, which increased the relative sink demand per leaf, although leaf export and stem sugar transport speeds did not change. Subsequently, Vincent et al. (2025) showed that apical leaves accumulate more starch than basal leaves when root growth is the only sink, and eventually deplete nearly all starch reserves in support of new shoots growing at the apex. Given that flushing increased leaf C export, on the basis of the source–sink attenuation hypothesis, we expected mature citrus leaves to respond to new shoot growth with increased CO_2_ assimilation. However, Wallace (1954) reported that mature leaves lost 25–30% of soluble N to new shoot growth. Thus, while providing a strong C sink, new shoots may also be a strong N sink. Given that leaf position impacts C export and allocation in response to the strong C and N sink dynamics imposed by new shoot growth, we conducted a study to assess C fixation and transport dynamics in response to flushing.

We hypothesized that (1) the emergence and development of new shoot flushes increase sink demand, which in turn stimulates both C export and photosynthesis in mature source leaves, and (2) apical and basal leaves differ in their physiological responses, reflecting their distinct roles in C allocation. To address these hypotheses, we studied mature citrus leaves across the course of vegetative phenological cycles and used gas exchange to assess their photosynthetic capacity and activity; ^14^C labeling to address C transport; and various methods to assess N, chlorophyll, and Rubisco contents. The results offer new insights into how source leaves integrate internal sink signals to adjust C metabolism, thereby advancing our understanding of source–sink regulation in citrus and other evergreen woody perennials.

## Materials and methods

### Plant material and growth conditions

In November 2022, healthy sweet orange plants (*C. sinensis* [L.] Osbeck), cv. Valencia 1-14-19, which were grafted onto ‘US-942’ rootstock (*C. reticulata* ‘Sunki’ x *Poncirus trifoliata* ‘Flying Dragon’), were sourced from a commercial nursery. The plants were acclimated in a controlled greenhouse environment at the University of Florida, Citrus Research and Education Center in Lake Alfred, FL, USA (28.1021° N, 81.7121° W). The plants were subsequently transplanted into 10-L containers filled with a ‘Pro-mix BX’ mixture (Premier Tech Ltd., Quebec, Canada) to ensure uniform growth conditions. In February 2023, the plants underwent pruning to promote uniformity, with each plant pruned to maintain 4–5 evenly distributed branches along the main stem. New shoots began to emerge 7–10 days post-pruning, and the number of shoots was controlled for consistency across the plants. For experimentation, one branch per plant was selected, featuring 8–9 fully expanded leaves on mature flushes (Table S1) and measuring approximately 14–15 cm in length. Once mature, these branches were used for all measurements over the subsequent flushing cycle.

Throughout the experiment, the greenhouse temperature was maintained between 24°C and 35°C, with the relative humidity ranging from 64% to 76%. The temperature fluctuations were controlled by a pad and fan system, which was automatically triggered when the temperature reached 30°C. Fertilization was provided biweekly, with 2.5 g of 20-20-20 fertilizer applied to the surface of each pot. Weekly pest inspections were conducted following the guidelines outlined in the citrus management guide (Diepenbrock *et al*., 2019).

### Experimental Design

This study employed a three-stage experimental design to investigate the effects of phenological stage on C dynamics in sweet orange plants (Fig. S1). The three stages were as follows: mature branches only (stage 1), mature branches with the initiation of new shoot growth (stage 2), and mature branches with fully expanded new shoots (stage 3).

At each stage, four trees were selected for analysis. Gas exchange measurements and C labeling experiments were performed on both the apical and basal leaves of the same branch. All selected leaves were healthy and fully expanded, ensuring consistency across the experimental setup.

### Branch Length and Leaf Count

Branch length was measured using a tape measure, and the number of leaves on each branch was manually counted. These data were used to characterize the branches and ensure uniformity across the experimental stages.

### Gas exchange measurement

Gas exchange parameters were measured using an infrared gas analyzer (LI-6800, LI-COR Biosciences, Lincoln, NE, USA) to determine the net assimilation rate (*A*), transpiration rate (*E*), intercellular CO_2_ concentration (*C*_i_), and stomatal conductance (*g*_sw_). Measurements were conducted between 09:00 a.m. and 11:00 a.m. under steady light intensity (1200 µmol m^-2^ s^-1^) and a CO_2_ concentration of 400 µmol mol^-1^.

### Dynamic Assimilation CO_2_ Response Curves

Dynamic assimilation CO_2_ response curves were generated using the LI-6800, following the procedure described by Saathoff and Welles (2021). The *A*/*C*_i_ response curves, which assess net CO_2_ assimilation under saturating light to intercellular CO_2_ concentrations, were generated with the following parameters: a water vapor mole fraction of 25 mmol H_2_O mol^-1^, a photosynthetic photon flux density of 1400 µmol m^-2^ s^-1^, and a constant flow rate of 300 µmol s^-1^. The CO_2_ concentration started at 100 µmol mol^-1^ min^-1^ and increased at a rate of 100 µmol mol^-1^ min^-1^ until it reached 1400 µmol mol^-1^.

### Estimation of Photosynthetic Parameters

The raw RACiR data were processed using the racir package (Stinziano *et al*., 2019) to create a calibrated *A*/*C*_i_ curve. The function fitaci() from the plantecophys package (Duursma, 2015) was used to estimate photosynthetic parameters, including the *V*_cmax_, *J*_1400_, TPU, and two transition points: the transition from carboxylation to electron transport rate limitation states (*C*_itrans1_) and the transition from the electron transport rate to triose phosphate utilization limitation states (*C*_itrans2_). Additionally, stomatal limitation to photosynthesis was calculated by comparing the value of *A* for each leaf to what value of *A* would be if *C*_a_ = *C*_i_ (e.g., *C*_i_ = 400 ppm) based on the *A*/*C*_i_ response curves.

### Chlorophyll measurement

Leaf discs were placed in 2 mL Eppendorf tubes with 1 mL of DMSO and stored in the dark at room temperature. The next day, the solvent was transferred to a new tube, and another 1 mL of DMSO was added to the leaf discs. After 3–4 days, the pigments were fully extracted, turning the solvent green and the discs translucent. The solution was mixed, and 200 μL was pipetted into a well plate with DMSO as the blank. The absorbances at 665 nm, 649 nm, and 480 nm were measured using a spectrophotometer. The chlorophyll content was calculated via the following equation:

Chla+b [μg] = [(A_665_ - A_700_) * 8.02 + (A_648_ - A_700_) * 20.2] * S

Chla+b [μg g^-1^] = Chla+b [μg]/FW

where A_665_ is the absorbance at 665 nm, A_648_ is the absorbance at 648 nm, A_700_ is the absorbance at 700 nm, S is the amount of solvent used, and FW is the fresh weight of the sample.

### Nitrogen content analysis

The leaf samples were dried in a forced-air oven at 55–60°C for 72 hours to remove excess moisture. The dried samples were then ground into a fine powder using a ball mill to ensure a uniform particle size. Each powdered sample (100 mg) was subsequently analyzed on a Leco CN828 Carbon and Nitrogen Analyzer (Leco Corporation, St. Joseph, MI, USA) to determine the N content.

### Rubisco Quantification

The Rubisco content in the leaf tissues was measured using a competitive ELISA. Fresh leaf samples were ground in liquid nitrogen with a mortar and pestle, and proteins were subsequently extracted using the 1× sample extraction buffer supplied with the Rubisco ELISA Kit (Novus Biologicals, NBP2-60142) at a tissue-to-buffer ratio of 1:5 (w/v). The extracts were immediately stored at -80□°C until further analysis.

Quantification was performed according to the manufacturer’s instructions. Briefly, samples were loaded into 96-well microplates precoated with Rubisco antigen. In this competitive assay, the Rubisco sample competes with the immobilized antigen for binding to a specific antibody. HRP-conjugated goat anti-rabbit IgG was added for detection. Upon addition of the substrate, a blue color developed, which turned yellow after acid neutralization. The absorbance was measured at 450□nm with background correction at 540□nm using a FlexStation 3 spectrophotometer (Molecular Devices, USA). The final Rubisco concentrations were determined by comparing the sample absorbance to a standard curve. Seven calibration standards (0, 3.12, 6.25, 12.5, 25, 400, and 800□μg/mL) were prepared to generate a standard curve. The relationship between the absorbance and Rubisco concentration was modeled using a four-parameter logistic (4PL) regression equation. All measurements were performed in triplicate across three biological replicates. Negative control wells lacking samples, antibodies, and HRP conjugates were included to assess the background signal and ensure assay specificity.

### Statistical analysis

Statistical analysis was performed using Statistix 8.1 software (Analytical Software, Tallahassee, FL, USA). The data were subjected to analysis of variance (ANOVA) to determine the significance of differences among the treatments and stages. When significant differences were detected, the means were compared using the honestly significant difference (HSD) test at a significance level of *p*≤0.05. Correlation analysis was performed in R (version 4.2.2, https://www.R-project.org/), using the ggpairs() command in the {Ggally}package (Schloerke B *et al*., 2024), which includes Pearson correlations and their *P* values for the correlations among variables throughout the entire experiment and within the phenological stage or within branch position.

## Results

### Trend of Photosynthetic Capacity in Citrus Leaves

The developmental stage of the new flush (hereafter “stage”) had a highly significant effect on all measured photosynthetic parameters in the mature leaves except for *C*_itrans2_ (*p*=0.6271), whereas leaf position (apical vs. basal) influenced only TPU (*p*=0.0253) and *C*_itrans1_ (*p*=0.0068) (Fig. 1). Specifically, *V*_cmax_, *J*_1400_, and TPU all increased from stage 1 to stage 2 and then decreased from stage 2 to stage 3. *R*_d_ also exhibited a significant stage□×□position interaction: at stage□2, the basal leaves maintained a greater *R*_d_ than did the apical leaves. *C*_itrans1_ displayed a parallel interaction, with higher *C*_itrans1_ values in basal leaves than in apical leaves at stage□3 (Fig. 1).

**Fig. 1.**
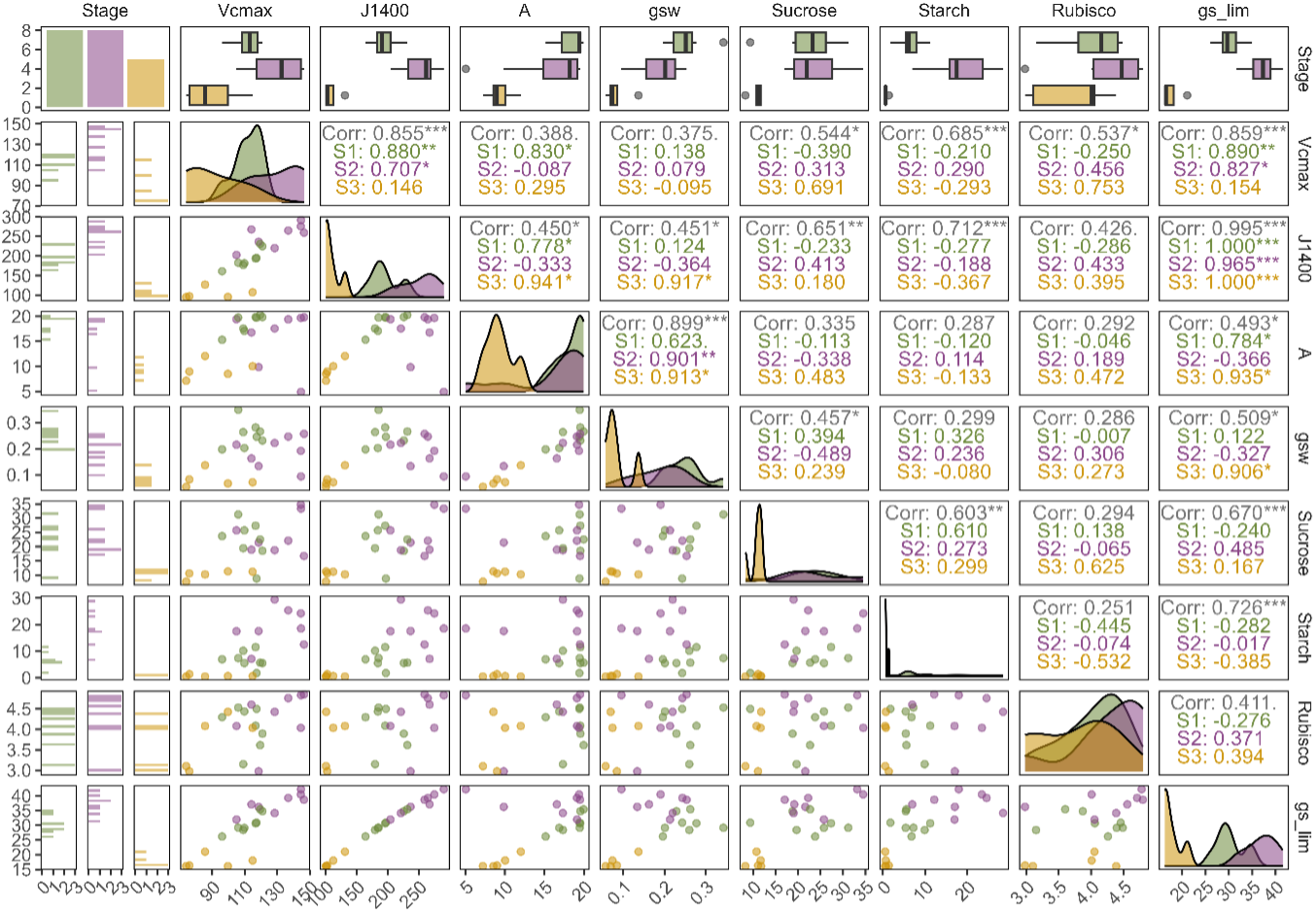
Comparison of photosynthetic parameters (*V*_cmax_, *J*_1400_, *TPU*, *C*_itrans1_, and *C*_itrans2_) in the apical and basal leaves of a branch at three phenological stages of sweet orange plants (*Citrus sinensis* [L.] Osbeck). P indicates the effect of the leaf position across the branch, S indicates the effect of the phenological stage, and PxS indicates the interaction between the leaf position and the phenological stage. The error bars denote standard errors, and the points denote means.

### Gas□Exchange Dynamics

Consistent with the changes in biochemical capacity, whole□leaf gas exchange parameters (*E*, *A*, *C*_i_, *g*_sw_ and *g*_sw_ limitation) were strongly influenced by stage (Fig. 2). In detail, at the ambient CO_2_ concentration, *E*, *A*, *C*_i_, and *g*_sw_ declined progressively from stage□1 through stage□3, reflecting reduced stomatal opening and CO□ uptake as the flush matured. Stomatal limitation (*g*_sw_ limitation) behaved differently: it rose from stage□1 to stage□2, indicating increasing diffusion constraints as new sinks emerged and then fell sharply by stage□3. Leaf position had no detectable main effect or interaction effect on these gas exchange traits.

**Fig. 2.**
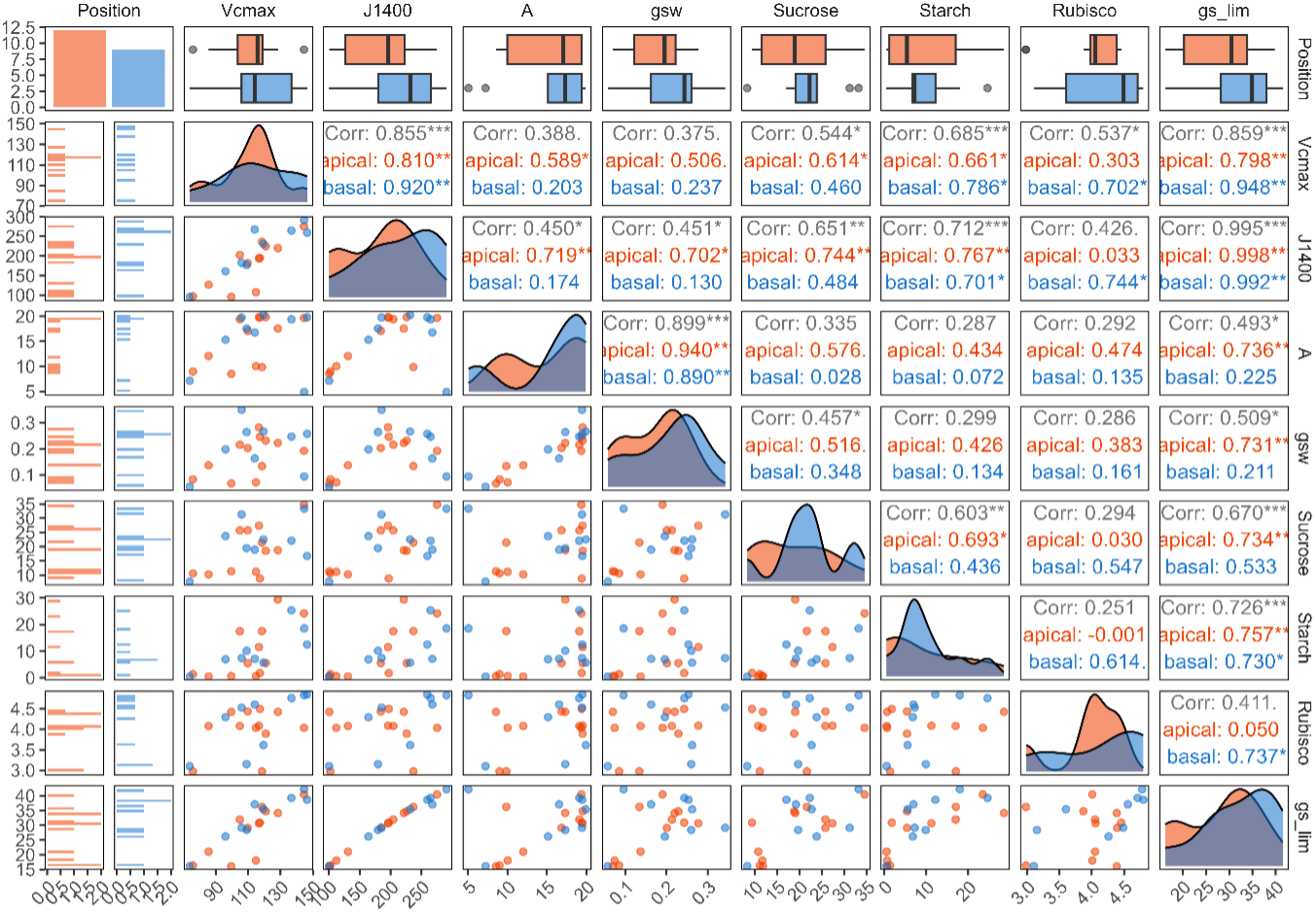
Comparison of gas exchange parameters (*E*, *A*, *C*_i_, *g*_sw_ and *g*_sw_ limitation) in the apical and basal leaves of a branch at three phenological stages of sweet orange plants (*Citrus sinensis* [L.] Osbeck). P indicates the effect of the leaf position across the branch, S indicates the effect of the phenological stage, and PxS indicates the interaction between the leaf position and the phenological stage. The error bars denote the standard error, and the points denote the means.

### C Export Patterns in Citrus Leaves

The whole□leaf ^14^C export rates were significantly affected by leaf position (*p*=0.0056; Fig. 3). Basal leaves exported more C at stage 2 than at stage 1 and stage 3, whereas apical leaves presented the opposite trend. Notably, at stage 2, the basal leaves exported approximately 82% of the newly fixed C, whereas 56% of the apical leaves did. No significant position differences were observed at stages□1 or□3, suggesting that positional partitioning of exported assimilates is most pronounced during the onset of new sink growth.

**Fig. 3.**
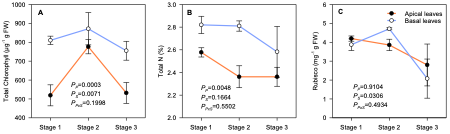
Comparison of the total chlorophyll, nitrogen and Rubisco contents in the apical and basal leaves of a branch at three phenological stages of sweet orange plants (*Citrus sinensis* [L.] Osbeck). P indicates the effect of the leaf position across the branch, S indicates the effect of the phenological stage, and PxS indicates the interaction between the leaf position and the phenological stage. The error bars denote standard errors, and the points denote means.

### Chlorophyll, Nitrogen, and Rubisco Contents

The total chlorophyll content mirrored this pattern, increasing from stage□1 to□2 and then decreasing at stage□3 (*p* = 0.0003), with basal leaves consistently maintaining higher pigment levels than apical leaves did (*p* = 0.007; Fig.□3A).

The leaf total N content was significantly influenced by the leaf position (*p* = 0.0048; Fig. 3B). Basal leaves contained more N than apical leaves did in stages□1 and□2, but basal N declined by ∼16% in stage□3, suggesting remobilization or dilution. The Rubisco content peaked at stage□2 (*p* = 0.0306; Fig. 3C), indicating elevated photosynthetic capacity during flush initiation, and then decreased by stage□3. Position did not influence Rubisco abundance.

### Correlations among Physiological Parameters

Correlation analyses revealed several consistent relationships among photosynthetic capacity, carbohydrate status, and stomatal behavior (Fig. 4). Across leaf positions, *V*_cmax_ was strongly correlated with *J*_1400_ and *g*_sw_ limitation, underscoring the coordinated regulation of carboxylation and electron transport with stomatal control. Importantly, *V*_cmax_ and *J*_1400_ were positively correlated with Rubisco in basal leaves but not in apical leaves, suggesting that Rubisco abundance may play a stronger role in determining carboxylation capacity in basal leaves. This finding is consistent with the greater fluctuations in *V*_cmax_ and Rubisco observed in basal leaves across stages. *A* was correlated with *g*_sw_ at both leaf positions, reflecting the central role of stomatal conductance in driving CO□ assimilation.

**Fig. 4.**
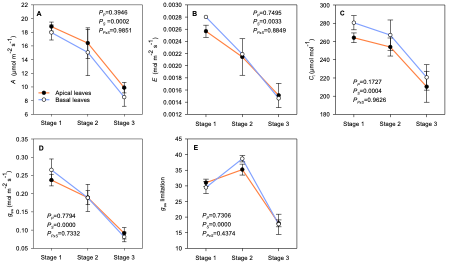
Correlation matrices among key physiological parameters in apical and basal citrus leaves. The variables shown include the maximum rate of Rubisco carboxylase activity (*V*_cmax_), the maximum rate of photosynthetic electron transport under saturating light (*J*_1400_), net photosynthesis (*A*), stomatal conductance (*g*_sw_), stomatal limitation (*g*_s_-lim), and sucrose, starch, and Rubisco contents.

Stagewise analyses (Fig. 5) further supported these relationships. *V*_cmax_ and J1400 were consistently correlated with *g*_sw_ limitation at stages 1 and 2, whereas at stage 2, both traits were also correlated with Rubisco. *A* maintained a strong positive association with *g*_sw_ at later stages, indicating that stomatal regulation continued to constrain assimilation as the flush progressed. Together, these patterns highlight how photosynthetic capacity and stomatal behavior are interdependent, with basal leaves showing stronger coupling of Rubisco to *V*_cmax_, which is consistent with their role in nitrogen mobilization during flush development.

**Fig. 5.**
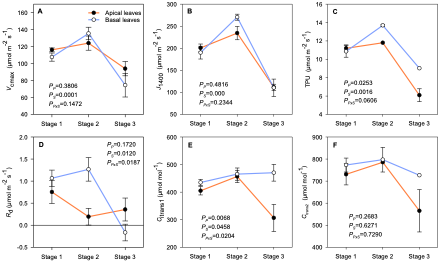
Stagewise correlation matrices among selected physiological traits in citrus leaves. The variables shown include the maximum rate of Rubisco carboxylase activity (*V*_cmax_), the maximum rate of photosynthetic electron transport under saturating light (*J*_1400_), net photosynthesis (*A*), stomatal conductance (*g*_sw_), stomatal limitation (*g*_s_-lim), and sucrose, starch, and Rubisco contents.

## Discussion

### Sink Demand Does Not Enhance Photosynthesis in Mature Leaves

We hypothesized that the emergence of a new vegetative flush would increase sink demand, leading to greater C export and an increase in photosynthesis in mature source leaves, which is consistent with the source–sink attenuation hypothesis. However, our results revealed a different story. Although C export increased during flush initiation (stage 2; detailed in (Vincent *et al*., 2025), photosynthetic activity (*A*, *g*_sw_) and photosynthetic capacity (*V*_cmax_, *J*_1400_) decreased from stage 1 to stage 3. This response contradicts the source–sink attenuation model, which predicts stimulation of photosynthesis under elevated export demand. Instead, our findings suggest that flush-induced sink demand exceeds the capacity of mature leaves to sustain C assimilation, ultimately leading to depletion of leaf carbohydrate reserves, as reported previously (Vincent *et al*., 2025).

The decline in *A* despite increased C exports reflects a coordinated, stage-dependent shift in photosynthetic limitation. During early flush initiation (stage 2), reduced *A* was driven primarily by a decline in *g*_sw_, whereas during later flush development (stage 3), further reductions in *A* were associated with decreases in *V*_cmax_ and *J*_1400_. These later declines coincided with reduced Rubisco abundance and modest changes in total N. Correlation analysis reinforced this interpretation, as *V*_cmax_ and *J*_1400_ were strongly associated with Rubisco content, particularly in basal leaves, indicating that the degradation of photosynthetic proteins directly constrained carboxylation capacity as sink demand intensified.

Importantly, this two-phase response, initial stomatal restriction followed by biochemical downregulation, aligns closely with the metabolic and proteomic reprogramming observed in our companion study (Chen et al., 2026, *Under Review*). This work demonstrated a broad downregulation of photosynthetic and primary metabolic pathways in mature leaves during flush development, alongside the mobilization of C and N reserves to support sink growth. Together, these findings suggest that mature citrus leaves undergo an active functional transition from C assimilation toward resource export and support rather than simply responding passively to carbohydrate feedback.

*g*_sw_ is known to be influenced by sucrose availability, which plays roles in both osmotic regulation and metabolic signaling within guard cells. In our companion study, sucrose levels declined sharply as sink demand increased (Vincent *et al*., 2025), whereas the present study revealed a positive relationship between *g*_sw_ and sucrose. Together, these findings suggest that sucrose limitation impairs stomatal function by restricting osmotic or metabolic signaling within guard cells, thereby increasing stomatal limitation during early flush growth. Similar sucrose-dependent regulation of the stomatal aperture via hexokinase-mediated signaling has been reported previously (Lima *et al*., 2018; Medeiros *et al*., 2018). Moreover, citrus girdling and sink manipulation experiments have shown that the photosynthetic rate can decrease before carbohydrate accumulation occurs, indicating early photosynthetic downregulation in response to export stress rather than classic feedback inhibition (Iglesias *et al*., 2002; Nebauer *et al*., 2011). Collectively, the integrated decline in *g*_sw_, *V*_cmax_, and Rubisco contents reflects a coordinated downregulation of photosynthesis under sustained sink demand, which is consistent with carbohydrate depletion and metabolic reprogramming (Vincent *et al*., 2025). These dynamics are distinct from those of natural leaf senescence, as the leaves examined here were young (2 and 3 months old) relative to the typical citrus leaf lifespan (approximately 17 months) (Wallace *et al*., 1954).

### Positional Differences in Export but not Photosynthetic Capacity

Although leaf position influenced export dynamics, it had a minimal effect on photosynthetic parameters. During flush initiation (stage 2), the basal leaves exported a greater proportion of newly fixed carbon than did the apical leaves (Vincent *et al*., 2025). This positional contrast is consistent with prior findings in citrus, where basal leaves are more strongly connected to structural sinks such as roots and stem tissues, whereas apical leaves preferentially serve local growing tips (Li *et al*., 2024). Despite these export differences, both the apical and basal leaves presented similar decreases in *A* and *g*_sw_ and increases in *g*_sw_ limitation across the phenological stages, indicating that positional differences in export did not translate into differential regulation of photosynthesis. This coordination of source activity within the branch occurs in response to the whole shoot sink demand.

The positive correlations of *V*_cmax_ and *J*_1400_ with Rubisco in basal leaves indicate stronger biochemical coupling between N investment and photosynthetic capacity in those leaves than in apical leaves. This relationship aligns with the greater variability in *V*_cmax_ and Rubisco observed in basal leaves, which is consistent with reports of preferential N remobilization from older or basal tissues (Xiong *et al*., 2025). In contrast, the apical leaves presented weaker correlations between Rubisco and *V*_cmax_, suggesting that their photosynthetic adjustments are less constrained by N-related biochemical limitations and more influenced by local sink connectivity. Together, these findings indicate that while export is not strictly determined by source capacity, the mechanisms supporting C allocation may differ by leaf position: basal leaves function as long-distance suppliers linked to protein and N turnover, and apical leaves serve emerging shoot sinks through carbohydrate-driven supply. This spatial partitioning of function aligns with recent work by Zhao *et al*. (2025), who showed that C allocation in citrus leaves is shaped by genotype and leaf type, likely through transcriptional regulation of sugar metabolism and transport genes. While transcriptomic data were not collected in the present study, our physiological findings support a model in which export is differentially regulated by leaf position, superimposed on a shared pattern of declining assimilation under increasing sink demand.

### Nitrogen Allocation: Stability with Functional Retooling

N allocation may play a major role in the photosynthetic responses observed in the present study. Although new vegetative flushes are strong C sinks (Li and Vincent, 2022), they are also major N sinks that require rapid acquisition of amino acids and proteins to support leaf expansion and metabolic activity (Wallace *et al*., 1954; Xiong *et al*., 2025). A recent proteomic study by Xiong *et al*. (2024) revealed that citrus trees remobilize N from stems and leaves to support new shoot growth. Under high N supply, stem reserves are the primary source of new shoot N, whereas under low N supply, leaves degrade proteins, including Rubisco and vegetative storage proteins (VSPs), to fuel new flushes. In the present study, total leaf N remained relatively stable across phenological stages, indicating that mature leaves did not experience wholesale N depletion. However, the Rubisco content decreased markedly by stage 3, which coincided with reductions in *V*_cmax_ and *J*_1400_, suggesting that N was selectively reallocated away from the photosynthetic machinery. Correlation analysis supported this interpretation, as *V*_cmax_ and *J*_1400_ were correlated with Rubisco content, particularly in basal leaves. These relationships indicate that reductions in carboxylation capacity were driven primarily by Rubisco loss rather than by changes in overall N availability. The stronger coupling observed in basal leaves is consistent with their greater functional role in supporting long-distance sink demand and N mobilization.

This pattern closely aligns with the proteomic evidence reported in our companion study (Chen et al., 2026 *under review*), which demonstrated the selective degradation of photosynthetic proteins, including Rubisco, alongside the maintenance of total N pools. This study further revealed broader remodeling of leaf protein composition, characterized by downregulation of C fixation pathways and enrichment of proteins associated with N mobilization and transport. Together, these findings indicate that N remobilization during flush development occurs through targeted protein turnover rather than through generalized N loss.

Importantly, the temporal sequence observed here suggests a coordinated shift in the N use strategy. Stomatal limitation dominated early during flush initiation (stage 2), whereas Rubisco degradation and reduced carboxylation capacity became evident only as sink demand persisted into stage 3. This delay implies that N reallocation from photosynthetic proteins is not an immediate response to sink activation but rather a regulated adjustment once sustained sink dominance is established. Such a strategy would allow mature leaves to maintain short-term assimilation while progressively prioritizing nitrogen export to developing shoots.

Although total leaf N changed only slightly, the functional consequences of N redistribution were substantial. The uncoupling of photosynthetic capacity from total N content reflects a shift in N partitioning away from C assimilation and toward supporting sink growth. Similar patterns have been reported in evergreen species, where the selective degradation of Rubisco and other photosynthetic proteins enables N conservation while reallocating resources to developing tissues (Xiong *et al*., 2025). In this context, the decline in Rubisco and *V*_cmax_ observed here represents a strategic retooling of mature leaves under high sink demand rather than premature senescence or nutrient deficiency.

## Conclusion

This study demonstrated that vegetative flush initiation in citrus plants stimulates C export from mature leaves without increasing their photosynthetic capacity, providing a clear counterexample to the source–sink attenuation hypothesis. Instead, sustained sink demand for both C and N drives a coordinated downregulation of photosynthesis in mature leaves. Decreases in *A* and *g*_sw_ during early flush development reflect stomatal limitation associated with carbohydrate depletion, whereas later reductions in *V*_cmax_ and Rubisco abundance indicate selective protein retooling rather than wholesale N loss. Despite positional differences in export capacity, apical and basal leaves presented similar photosynthetic responses, underscoring the systemic coordination of source–sink interactions within branches. Together with complementary physiological, metabolic, and proteomic evidence (Vincent *et al*., 2025; Chen et al., 2026 *under review*), these findings support a model in which mature citrus leaves function as regulated C and N conduits under strong sink demand, prioritizing the establishment of new photosynthetic capacity over the maintenance of existing assimilation.

## Supplementary data

**Fig. S1.** Illustration of the research material. The top, orange- and blue-colored leaves indicate the ^14^C-labeled apical and basal leaves, respectively.

**Table S1.** Growth characteristics of the experimental trees.

## Author contributions

SBH, YW, and CV planned and designed the research. SBH, QM, and SL conducted the experiments. CV, SBH, and QM processed the data. SBH performed the statistical analysis. CV and SBH drafted the manuscript.

## Data availability

The data that support the findings of this study are available from the corresponding author upon reasonable request.

